# An expansive animal gut microbiome dataset elucidates major compositional shifts across bilaterian evolution

**DOI:** 10.64898/2026.04.29.721755

**Authors:** Samuel Degregori, Lucas Patel, Luis Xu, Michael Iter, Yuhan Weng, Isabella Huang, Harrison J. Martel, Dmitry Kisselev, Jianshu Zhao, Celeste Allaband, Gail Ackermann, Antonio González, Daniel McDonald, Katherine R. Amato, Rob Knight

## Abstract

Animal gut microbiomes provide key physiological functions and are critical for host health. They vary dramatically across the animal kingdom, and are shaped by factors including host diet, evolutionary history and environment. However, analyses of gut microbiomes spanning the entire metazoan clade are lacking, limiting our understanding of the fundamental principles governing gut microbiomes. Here we present the Gut Microbiome Tree of Life (GMToL), a curated 16S amplicon dataset of 17,366 samples from 1,553 host species across 26 host classes from 284 studies, enabling analysis of large-scale evolutionary trends. Using ancestral state reconstruction, we provide a critical baseline calculation of major compositional shifts in gut microbiomes throughout animal evolution. We show that the ancestral animal gut was likely dominated by Pseudomonadota. A major shift to Bacteroidota occurred during the evolution of tetrapods, followed by the emergence of Bacillota-dominated guts in mammals and birds. We identify conserved core gut microbes and demonstrate how GMToL can be leveraged to contextualize the evolutionary history of specific microbial taxa. Ultimately, this framework enables the predictive mapping of microbial symbionts across uncharacterized host lineages, and establishes a quantitative baseline for comparative microbiome research at scale.

## Introduction

Gut microbiomes are critical components of animal health and survival^1–3^. In the wild, all vertebrates have gut microbiomes that vary with host phylogeny^4,5^, diet^6–8^, and environment^9–11^. Animal gut microbiomes are therefore integral to understanding animal ecology and evolution.

Of the many host factors that have been identified as shaping gut microbiome diversity, host phylogeny (evolutionary history) and host diet have been identified as two of the most prominent in explaining gut microbiome variability across the animal kingdom. In mammals, for example, host phylogeny correlates strongly with gut microbiome variation^12,13^, particularly when compared to other host clades^5,14^. Diet can have a larger effect than host phylogeny, but the correlations between these two variables make them difficult to tease apart^12,15^. Effects of host diet within individual clades have been demonstrated in large-scale comparative datasets that can account for correlations between diet and phylogeny^8,16,17^, but how host diet shapes gut microbiomes across the entire animal kingdom is not yet understood.

Gut microbiomes are also influenced by the host’s external environment, but the magnitude of this effect varies across clades. In mammals, the environment plays a small role in shaping gut microbiome variation across host species^4,5^ and a larger role in shaping intra-species variation^11^. In other vertebrate hosts, the environment plays a larger role in shaping gut microbiome diversity, exemplified in fish^9,10^, amphibians^18^, and lizards^19,20^. However, whether these patterns extend to invertebrates is largely unknown. A comparative analysis of ant gut microbiomes^21^ found that habitat significantly impacts ant gut microbiomes, but efforts to test these findings across other insects and invertebrates at large have been limited^22^.

Several large-scale efforts have sampled gut microbiome diversity across a broad range of hosts. Youngblut et al. (2019) and Song et al. (2020) both provided diverse datasets of vertebrate gut microbiomes, which have served as a useful resource for further comparative studies ^16,23,24^. Analyses integrating vertebrates with other metazoan clades have been limited. The Animal Gut Microbiome Database (AMDB) includes 34 gut microbiome studies across 9 host classes of vertebrates and invertebrates totaling 2530 gut microbiome samples^25^. Ma et al. (2021) compiled 4903 animal gut microbiome samples across 10 host classes of vertebrates and invertebrates. However, more expansive efforts are timely and applying evolutionary tools such as ancestral state reconstruction^5,7,14^ across both vertebrate and invertebrate hosts can reveal deep evolutionary patterns spanning the animal kingdom. Moreover, including data from both comparative and single-host studies can dramatically improve host diversity in data aggregation efforts^16,26^. Many ecologically focused gut microbiome studies in individual host species yield publicly available data not included in comparative efforts.

Here, we leverage the largest compiled animal gut microbiome dataset to date (17,366 samples; 1,553 host species; 26 host classes), which includes an unprecedented number of underrepresented host taxa, to better understand how animal ecology and evolution shape gut microbiome diversity (Fig 1).

**Figure 1.**
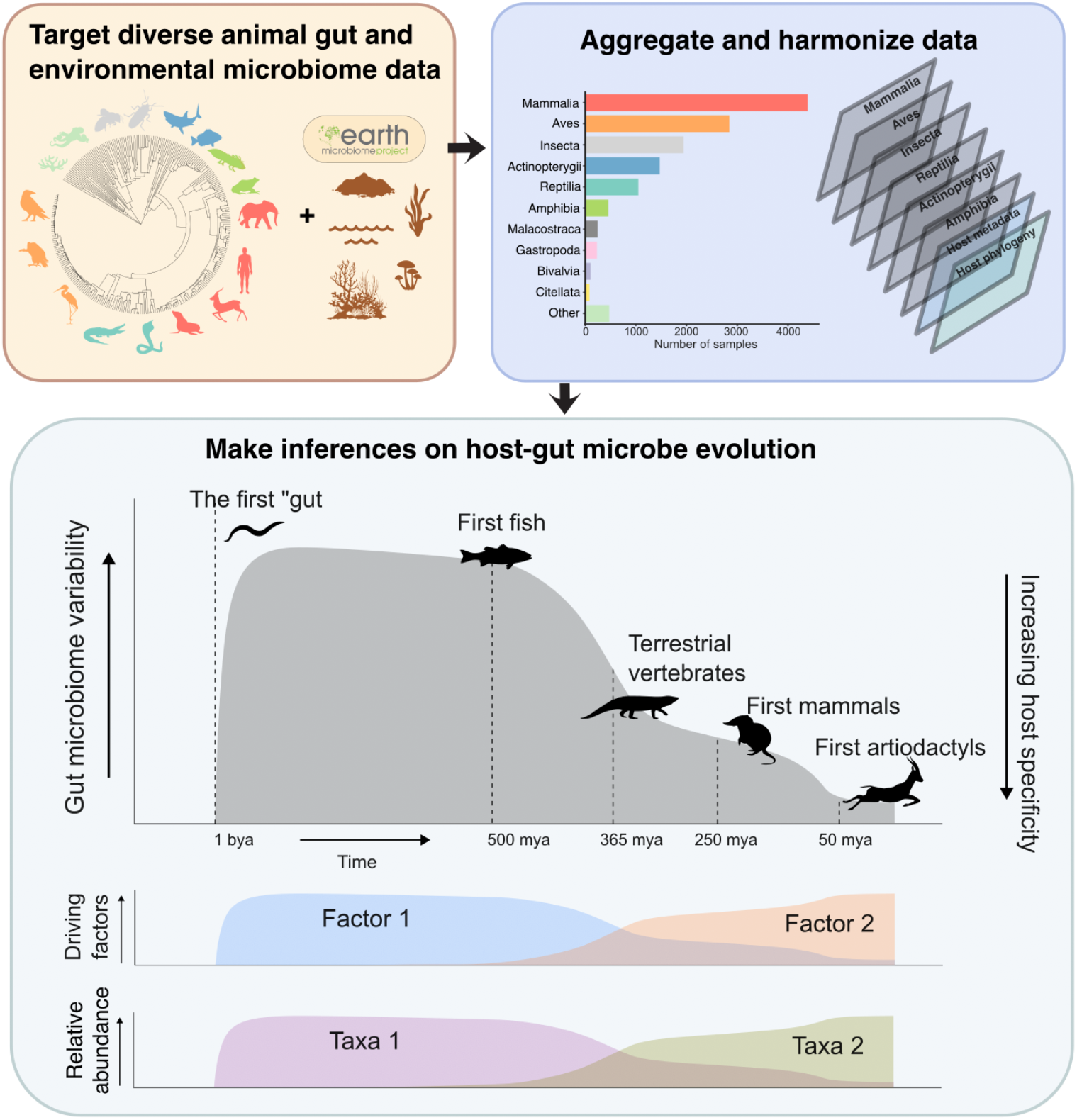
Workflow showing data collection, analysis, and synthesis.

## Results

## Data aggregation

Aggregating 284 animal gut microbiome studies across the tree of life resulted in 4,011,620 unique amplicon sequence variants (ASVs) totaling 694 million reads, averaging 29,553 reads per sample across 17,366 samples (see Table S1 for comparisons to previous aggregation efforts). After performing standardized global quality control by filtering out ASVs with a single read, ASVs present in only a single sample, and ASVs assigned to eukaryotes, chloroplasts, and mitochondria, we were left with 3,376,978 ASVs with a median of 27,744 reads per sample.

## Bacillota, Bacteroidota, and Pseudomonadota taxa dominate animal gut microbiomes and drive vertebrate and invertebrate differences

Three main phyla—Bacillota (Firmicutes), Bacteroidota (Bacteroidetes), and Pseudomonadota (Proteobacteria)—comprise most of the bacterial diversity across the animal gut microbiomes we aggregated (Fig 2A). Across the 26 host classes, all three phyla were present at 1% relative abundance or greater, but their proportions varied significantly and reproducibly across host phylogeny. For example, on average, 4.1% of Clitellata (worm) gut microbiomes consist of Bacillota, in striking contrast to the average of 51.5% in Mammalia. This trend held across host class with vertebrate classes averaging significantly higher proportions of Bacillota (36.5%, of which 24.6% consisted of the subphylum Bacillota_A, primarily of the class Clostridia) and 19.2% Pseudomonadota. This compared to invertebrates averaging 18.7% Bacillota (with only 3.8% consisting of Bacillota_A) and significantly higher proportions of Pseudomonadota (42.6%; see Table S2 for Kruskal-Wallis results for each of these comparisons). Terrestrial vertebrates, in particular, possessed consistent amounts of Bacillota, never dropping below 28%, but invertebrate classes did not exceed 21% Bacillota. Invertebrate gut microbiomes were also more similar in composition to the environmental samples than vertebrate gut microbiomes were (P_Mann- Whitney_<0.001). When considering within-group variation, vertebrate gut microbiomes were more similar to one another than to invertebrate gut microbiomes (P_Mann-Whitney_<0.001). The same did not hold true for within-invertebrate diversity compared to vertebrates.

**Figure 2.**
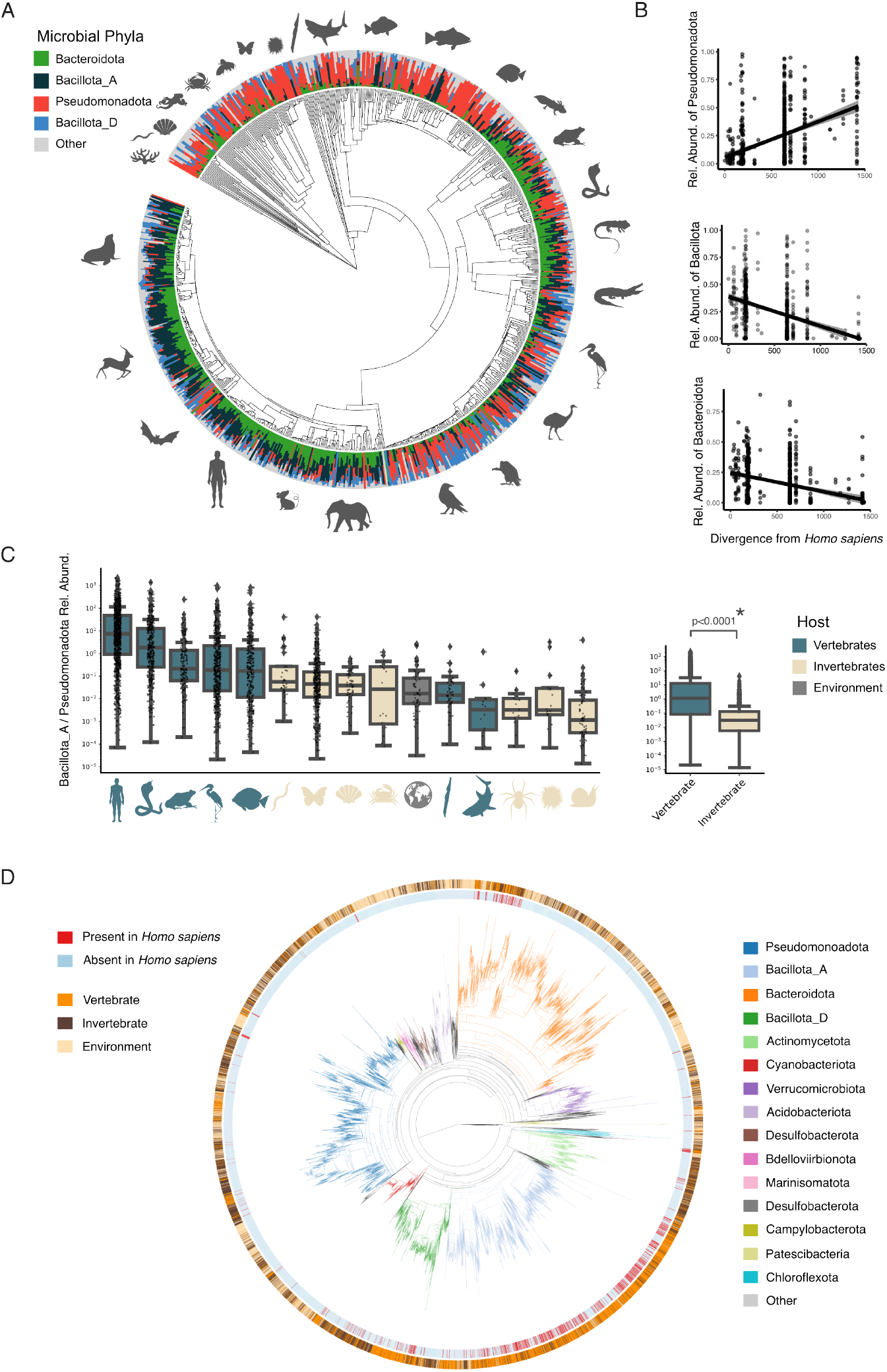
A) Gut microbiome composition across host phylogeny. Samples are merged by host species (n=828) that matched to NCBI. Stacked barplots at tips denote the relative abundance of major bacterial phyla. B) Relative abundance of Bacillota (including all subclades), Bacteroidota, and Pseudomonadota as a function of host divergence time from *Homo sapiens*. Host divergence was calculated based on the time- calibrated branch lengths of the host tree. C) The ratio of Bacillota_A *to* Pseudomonadota clr-transformed relative abundances across host classes with colors denoting vertebrates, invertebrates, or environmental samples and then the same data plotted across vertebrates and invertebrates on the plot to the right. D) A phylogeny of all identified gut microbes in our dataset that mapped to Greengenes2^27^ (74,384 ASVs) . The inside ring denotes whether ASVs were present in human hosts (red) or absent (light blue). The outer ring denotes whether ASVs were present in vertebrates (orange), invertebrates (brown), or environmental samples (beige).

When considering differences relative to the human gut microbiome (whereby all samples were merged at the host species level), we found that the relative abundance of Pseudomonadota (r=0.439, p<0.001, Fig. 2B) increased as host divergence time from humans increased, and the opposite trend held for Bacillota (r=-0.394, p<0.001) and Bacteroidota (r=-0.318, p<0.001). We found that the ratio of Bacillota_A to Pseudomonadota (Bac:Pseud) varied significantly across classes, with terrestrial vertebrates and fishes exhibiting the highest ratios (Fig 2C) and vertebrates overall exhibiting significantly higher ratios than invertebrates (H=582, p<0.001). When filtering our entire dataset against the Greengenes2 database in its state prior to this study^27^, we are left with 74,384 unique microbial ASVs (Fig. 2D). For comparison, the human samples alone comprised 1,052 microbial ASVs. In total, 28.0% of the identified animal gut ASVs belonged to Bacillota, 25.1% belonged to Pseudomonadota and 18.6% belonged to Bacteroidota. The new ASVs have been incorporated into a new release of Greengenes2 that expands its utility for future studies of this type.

Animal gut microbiome composition varied significantly across host class (F=43.15, P_PERMANOVA_=0.001, df=25, Fig 3A). Terrestrial vertebrates split into two prominent clusters: one consisting of birds, bats, and snakes, and the other consisting of mammals, non-snake reptiles, and amphibians (F=279, P_PERMANOVA_=0.001, df=2, Fig 3B). The third cluster consisted of fish, all invertebrates, and environmental samples, and aligned more closely with birds, bats, and snakes, than with other vertebrates (P_Mann-Whitney_=0.001, Fig 3B). Host classes also differed significantly in alpha diversity (H=804.9, p=0.0001, df=24, Fig 3C), with Clitellata (worms), Merostomata (horeshoe crabs), and Chondrichthyes (sharks), in particular, having significantly higher alpha diversities than the rest of the animal kingdom: these classes had similar diversities to our reference panel of environmental samples (Fig 3C). Overall, 110 of the 325 pairwise comparisons between host alpha diversities were significantly different (Table S3). A heatmap analysis of the most abundant gut microbes across terrestrial vertebrates showed that an *Enterobacteriaceae_A sp*. was highly prevalent in all vertebrate groups except for fish and mammals and *Lachnospiraceae sp*. which was similarly prevalent in all vertebrate groups except fish or bats (Fig 3D).

**Figure 3.**
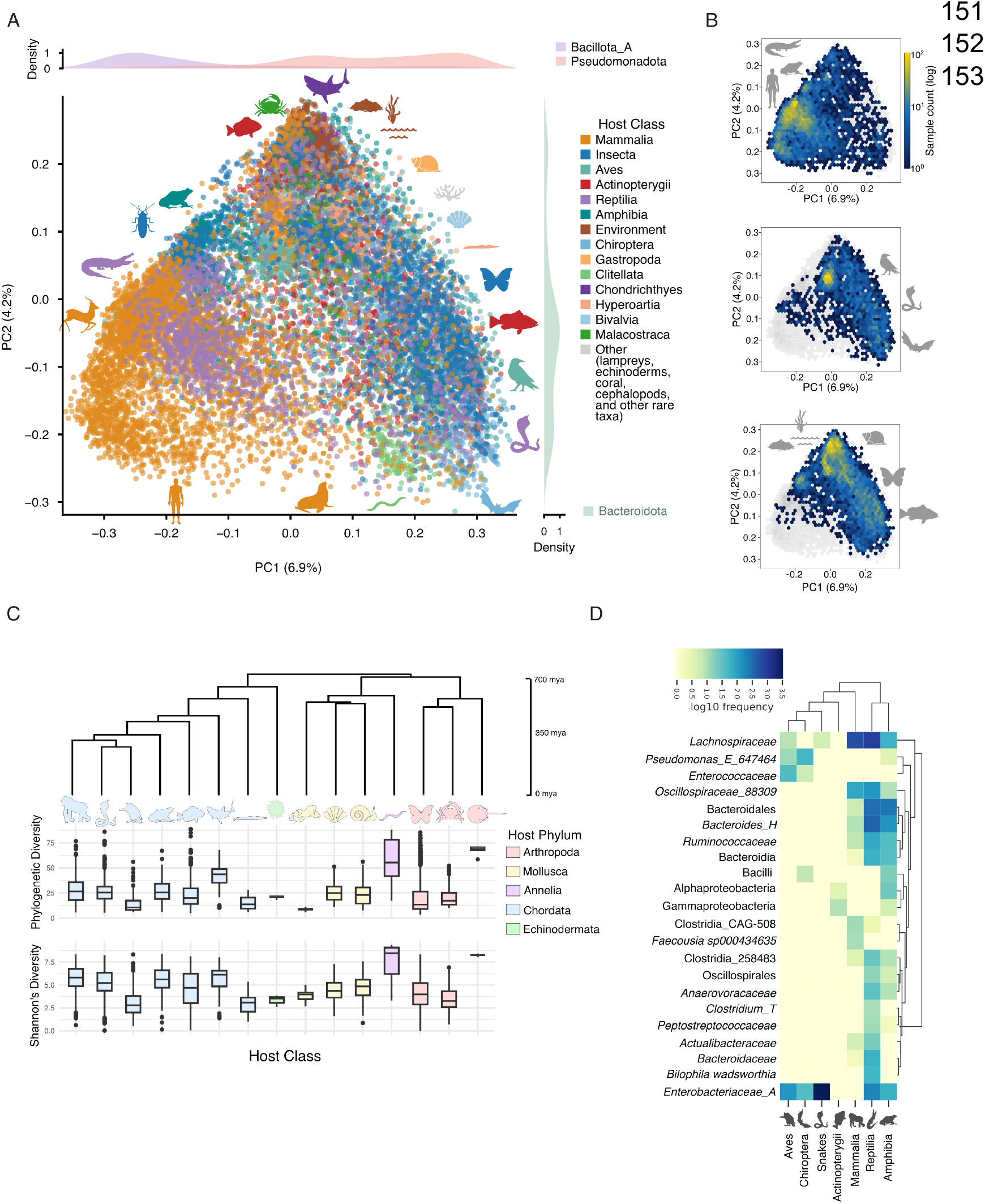
A) A PCoA plot based on unweighted UniFrac distances of animal gut microbiome spanning the tree of life (n=16,396). Colors denote host Class, except brown which denotes environmental microbiomes. Silhouettes serve as representative host taxa with the corresponding host Class color. For visualization purposes, not all host classes are colored, and instead binned to “Other”. B) Hexagonal bin plots depicting 3 significant clusters we identified (top plot: mammals, amphibians, and reptiles, excluding snakes; middle plot: birds, bats, and snakes; bottom plot all remaining classes grouped with environmental samples). C) Both Faith’s phylogenetic diversity and Shannon’s diversity across host classes. D) A heatmap of the relative abundance of prominent microbial taxa across vertebrate classes with two groups, snakes and birds, shown separately to highlight a convergence with birds.

### Ancestral state reconstruction infers four dominant ancestral clades of gut microbes and establishes baseline shifts throughout animal evolution

To infer ancestral states of host gut microbiome composition across the animal kingdom, we employed a multivariate Brownian Motion model^28,29^ using a time-calibrated host phylogeny tree we constructed using the Timetree of Life^30^. To identify major shifts in composition throughout animal evolution we mapped the most dominant ancestral microbial taxa, derived from our reconstruction analysis, across the host tree (Fig. 4) at the microbial Phyla and Order ranks. Our hosts of interest all fell within the Bilateria clade, spanning 23 Classes and 7 Phyla, including invertebrates with primitive guts such as worms (Annelida), brittle stars (Echinodermata), and mollusks (Mollusca). Our time-calibrated host tree placed the MRCA of our dataset at 627-830 million years ago (mya) and our reconstruction analysis infers Pseudomonadota as the dominant gut phyla throughout this time period. For fish and invertebrates specifically, Pseudomonadota stayed dominant up until the present. At an estimated 348-355 mya, markedly around the time of tetrapod emergence, a shift from Pseudomonadota to Bacteroidota occurs, suggesting the beginning of non-Pseudomonadota-dominated gut microbiomes. Another major shift occurs at an estimated 316-322 mya, marking amniote evolution, where amniotes split into a Bacillota_A- dominated clade consisting of all the mammalian hosts in the dataset. Amphibians and reptiles, apart from snakes, appear to retain Bacteroidota as their dominant microbial phyla, and the evolution of birds marks the introduction of Bacillota_D as a dominant clade at an estimated 97- 118 mya. At the Order level, the Orders Enterobacterales_A (Pseudomonadota), Bacteroidales (Bacteroidota), Clostridiales (Bacillota_A) and Lactobacillales (Bacillota_D) followed a similar pattern to their respective phyla, with the exception of mammals retaining Bacteroidales and only a small clade of carnivores shifting to Clostridiales. Thus, our analysis establishes a baseline estimation, spanning vertebrates and invertebrates, of the timing of major shifts in gut microbiome composition throughout animal evolution.

**Figure 4.**
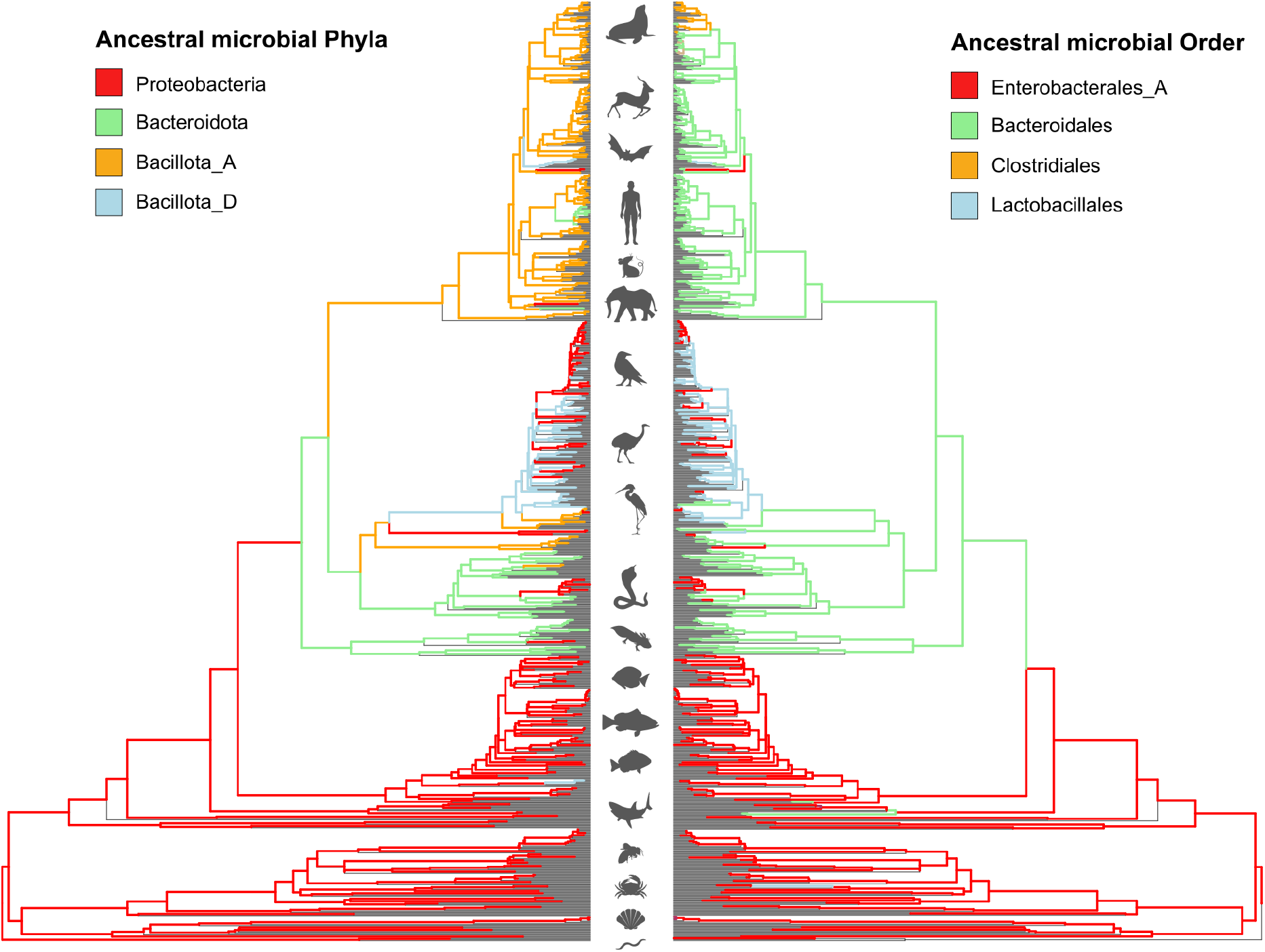
Ancestral reconstruction of dominant gut microbial phyla (left) and orders (right) across the host tree of life using ABDOMEN (v1.0). Treating gut microbiome composition as a trait, branch colors represent the highest relative abundance ancestral taxa calculated at each internal node throughout the tree. Host taxonomy for both trees is depicted by the host silhouettes at the tree tips.

### GMToL allows for identification of conserved gut microbes and provides evolutionary context for gut microbes of interest

Animal gut microbiomes as a whole shared a small but diverse range of taxa that comprised the core community across the animal kingdom (Fig 5A). The most prevalent gut microbe (defined solely by presence/absence and greater than 0.01% relative abundance of the ASV) across host classes was an undefined *Escherichia sp*. and *Corynebacterium kefirresidentiae* that were present in 20 of the 23 host classes considered and only absent in lobe-finned fish (Sarcopterygii), sea cucumbers (Holothuriida), brittle stars (Ophiuroidea) (Fig. S2). The second-most prevalent consisted of *Acinetobacter sp*., *Caldora sp010672925*, and *Lawsonella sp*. present in 18 host classes, and were also consistently absent from brittle stars and sea cucumbers, but not lobe-finned fish. Other prominent taxa included *Enterobacter, Phocaeicola_A, Streptococcus, Methylobacterium, Ralstonia*, and *Paraclostridium*. We then cross-referenced these taxa at the genus level against known contaminants that make up the “kitome”^31^. However, many of the taxa that overlapped with kitome genera were from the Pseudomonadota phyla. In fact, of the top 23 most prevalent microbes, 12 of them were Pseudomonadota, and 9 of these overlapped with known kitome genera. However, we cautiously include these taxa in our analysis because many of our animal gut microbiomes include a large range of common environmental microbes that are likely relevant to ectotherms and invertebrates, which have less barriers to host translocation. When considering microbes that were pervasive at the sample level, we did not find any ASVs that served as a core community across all individual samples even when a threshold of 50% prevalence or above was applied. We include a full list of prevalent taxa and the number of host classes they span in Table S4. Future studies using methods with higher taxonomic resolution will be important for resolving this issue definitively.

**Figure 5.**
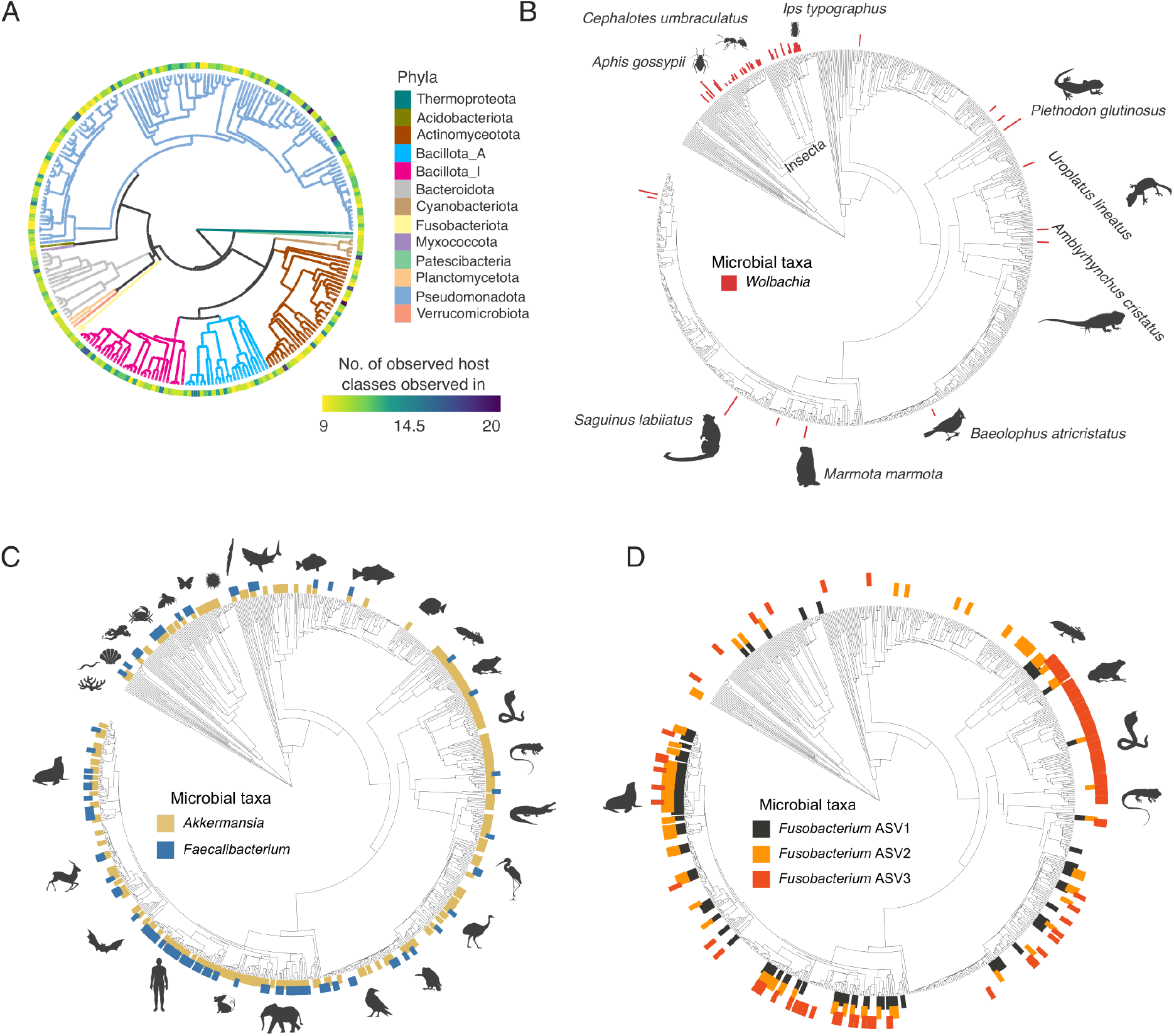
A) A microbial phylogeny of the most prevalent gut microbes (present in at least 9 host classes) across the entire dataset. B) A host phylogeny annotated by gut microbe *Wolbachia* relative abundance at the leaves. C) Host tree annotation of gut microbes of interest, including *Akkermansia* and *Faecalibacterium* and D) *Fusobacterium spp*. delineated by ASV number to avoid making inferences on species taxonomic assignment with solely 16S data. Here, microbes are considered present if greater than 0.01% relative abundance for a given host taxa.

We also show that GMToL can provide valuable host evolutionary context for gut microbes of interest. *Wolbachia*, a well-studied parasitic gut microbe in insect species^32^, is highly concentrated across a majority of insect hosts (Fig. 5B). *Akkermansia muciniphila*, a well-studied gut microbe in humans^33^, is quite prevalent across the host tree of life, while *Faecalibacterium prausnitzii*, also well-studied in human guts^34,35^, is largely confined to mammals and some marine invertebrates (Fig. 5C). Because *Fusobacterium* has been consistently implicated in colorectal cancer^36,37^, we investigated its prevalence across animal gut microbiomes. We found a *Fusobacterium_B sp*. and *Fusobacterium perfoetens* diversity to concentrate in mammalian carnivores while *F. ulcerans* was highly prevalent in amphibians and reptiles (Fig 5D).

### Animal host classes exhibit distinct gut microbiome composition and diversity

To identify uniquely enriched microbial taxa across each host class we employed a Bayesian inference approach specifically designed for sparse compositional data^38^. We found significantly enriched taxa across 7 host classes, spanning both vertebrate and invertebrate hosts, and spanning 33 microbial phyla but with more than 90% of the identified taxa comprising 10 phyla (Fig. 6A; Table S5). In fish, the most enriched gut microbe (sorted by mean estimates, *μ*) was an undescribed *Burkholderiales sp*. in the Pseudomonadota phylum (*μ* =15.1; CI: 12.0,18.6). While only 91 of the 542 (16.8%) enriched taxa we identified in fish were in the Pseudomonadota phylum, Pseudomonadota made up 50% of the top 20 and 80% of the top 10 enriched taxa in fish. Our Bayesian model also identified taxa in the Deinococcota phylum (extremophilic microbes) in the top 20 enriched taxa, including *Thermus* (*μ*=13.2; CI: 10.1, 16.6), *Deinococcus* (*μ*=12.8; CI: 9.7, 16.3), and *Meiothermus* (*μ*=12.7; CI: 7.1, 16.6). In mammals, out of the 860 enriched gut microbes our Bayesian model identified, 324 (37.7%) were Bacillota_A and 118 (13.7%) were Bacillota_D. All of the identified Bacillota_A taxa were *Clostridia spp*. The most enriched gut microbe in mammals was an undescribed *Christensenellales sp*. in the Bacillota_A phylum (μ=11.2, CI: 8.1, 14.8). Bacteroidota made up 14% of the enriched mammalian gut microbes including *Prevotella* in the top 10 (μ=10.9; CI: 7.0, 14.9). Mammalian gut microbiomes were also enriched with Archaea, notably *Methanobrevibacter* (*μ*=10.6; CI: 6.4, 14.5) in the top 20 of all enriched taxa in mammals. These Archaea taxa were also enriched in birds and even in fish (*μ*=4.8; CI: 0.6, 9.1). In insects, the top 20 enriched gut microbes were not dominated by a single phylum and included Pseudomonadota, Bacteroidota, Bacillota_A, Bacillota_D, Actinobacteriota, and Synergistota, and included the well-studied *Wolbachia* (*μ*=11.3; CI: 2.3, 18.6) which was not present in any other host class. In birds, 110 of the 810 enriched taxa belonged to the Bacillota_D phylum, including 5 of the top 10 enriched taxa, notably *Catellicoccus* (*μ*=10.8; CI: 4.6, 16.3) and *Mesomycoplasma* (*μ*=10.6; CI: 6.1, 15.7). Similar to mammals, reptiles were largely characterized by *Bacillota_A* taxa, 157 of the 382 enriched taxa (41.1%), notably *Cloacibacillus* (*μ*=12.5; CI: 8.7, 16.1) and *Pelospora* (*μ*=11.9, CI: 8.2, 15.5). Unlike the other terrestrial vertebrates, a majority of the most enriched taxa in amphibians were Pseudomonadota taxa (28.9% of the 97 taxa), including the most enriched of all 97, *Berkiella_A sp*. (*μ*=12.3; CI: 8.9, 16.7).

**Fig 6.**
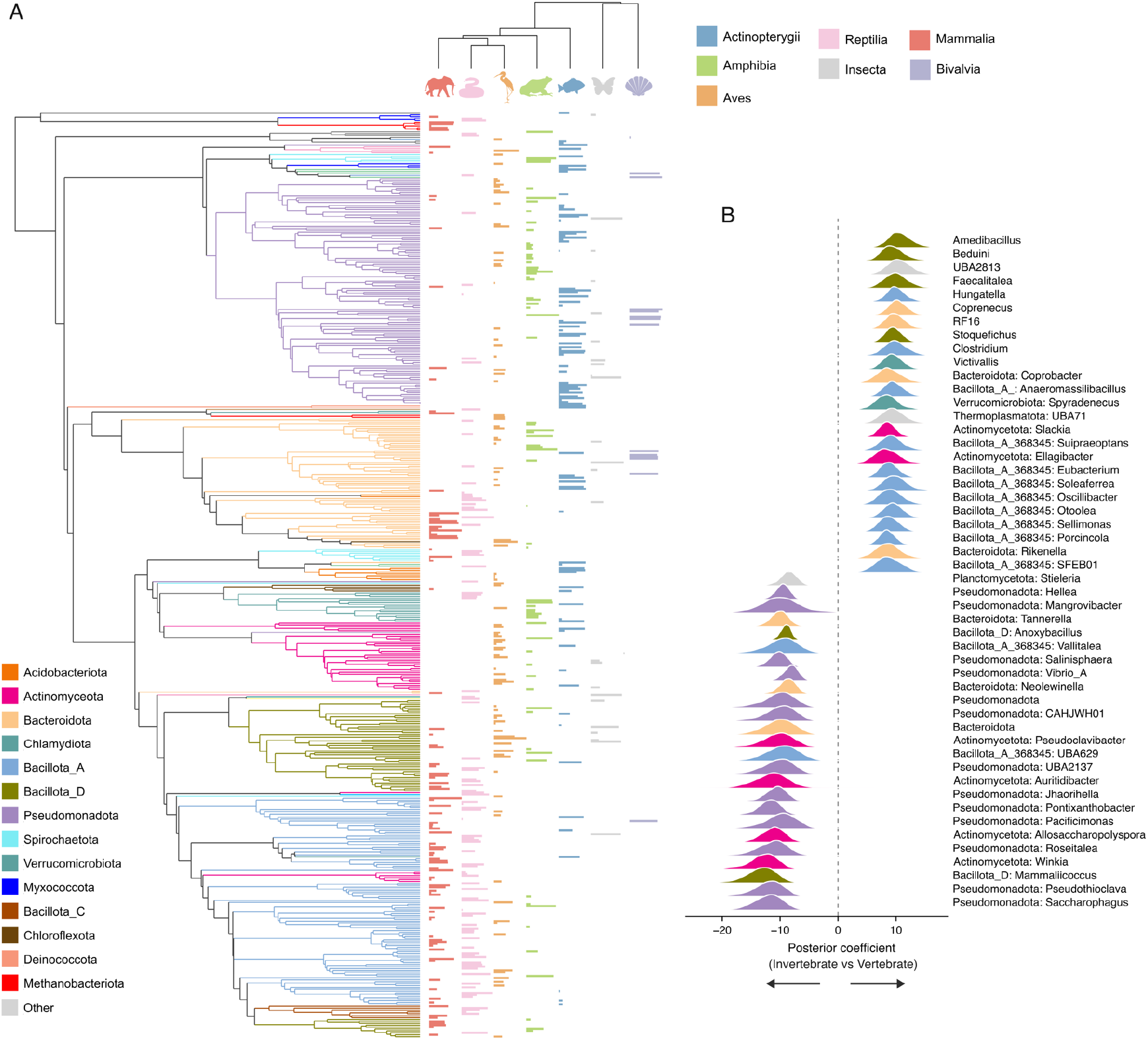
Differential abundance results displayed across A) a microbial phylogeny of the top most differentially abundant (“enriched”) taxa across 7 host classes. The phylogeny is colored by phyla and bars at the tips represent mean estimates of enriched taxa detected using BIRDMAn, a Bayesian inference model approach^38^. Each host class was compared against the rest of the dataset. For classes with more than 100 microbial taxa identified, we depict only the top 100 (sorted by mean estimates) for visualization purposes. B) Posterior distributions of the top 25 vertebrate-enriched (positive) and top 25 invertebrate-enriched (negative) genera identified by BIRDMAn. Each ridge represents the full posterior density from 2000 MCMC draws (4 chains × 500 iterations); the dashed vertical line marks zero (no difference). Curves are colored by microbial phylum for each microbial genus following the same color legend for (A) in the bottom left.

To determine differentially abundant genera across vertebrates and invertebrates, we employed BIRDMAn across these two groups. Vertebrate-enriched taxa spanned 46 phyla, dominated by Pseudomonadota (26.9%), Bacillota_A (22.7%), Bacillota_I (11.9%), and Bacteroidota (10.4%). Invertebrate-enriched taxa spanned a narrower 26 phyla but were disproportionately composed of Pseudomonadota (41.4%) and included several lineages with no cultured representatives, including candidate phyla CG2-30-53-67 and Eisenbacteria, highlighting the abundance of poorly characterized microbial diversity in invertebrate gut microbiomes. However, the well-studied *Wolbachia* (μ = ™1.64, CI: ™2.44, ™0.87), an obligate endosymbiont of arthropods, was significantly associated with invertebrate microbiomes, supporting our previous phylogenetic analysis of *Wolbachia*. Among the most strongly vertebrate-enriched genera were *Amedibacillus* (μ = 12.66, CI: 9.41, 15.91, Fig 6B), *Beduini* (μ = 12.54, CI: 9.30, 15.67), and *Faecalitalea* (μ = 12.20, CI: 8.70, 15.54), all within the Erysipelotrichaceae family. Vertebrate- enriched taxa also included the underscribed Cyanobacteria, *UBA2813* (μ = 12.44, CI: 9.17, 15.66), in the family Gastranaerophilaceae, an anaerobic cyanobacterial lineage known to colonize animal guts.

## Discussion

This study presents the most comprehensive aggregation of animal gut microbiomes to date, leveraging a dataset of over 17,000 samples across 284 datasets with 26 host classes represented. By integrating data from both large-scale comparative studies and previously separate single-species ecological research, we provide a high-resolution map of the “Gut Microbiome Tree of Life” within animal hosts. Our findings reveal that animal gut microbiomes are diverse, comprising more than 74,000 microbial taxa–far outnumbering the ∼1,000 taxa that are often found in humans. We also show that animal gut microbiomes follow generalizable patterns and provide valuable baseline measurements of ancestral gut microbiome composition and major compositional changes throughout animal evolution.

### Ancestral gut microbiome composition

Ancestral state reconstruction provides a temporal framework for understanding animal gut microbiome diversity in the context of host evolution. Our findings suggest that the “ancestral” animal gut was likely dominated by Pseudomonadota, a pattern that persists today in many marine invertebrates and fishes. The first major shift—a transition toward Bacteroidota dominance— coincides with the evolution of tetrapods approximately 350 million years ago. It remains unclear why Bacteroidota and Bacillota do not start to dominate gut microbiomes until tetrapod evolution. For example, previous work shows that fish gut microbiomes, particularly those of carnivores and herbivores, converge with mammalian gut microbiomes, largely due to Bacillota microbes^6^. However, our work here shows that despite this convergence, fish gut microbiomes are still largely dominated by Pseudomonadota. Thus, it may be that the evolution of more complex immune systems, paired with distinct changes in gastrointestinal architecture—such as increased gut retention times and the development of specialized crypt structures—in terrestrial vertebrates may have selectively facilitated the colonization for Bacillota and Bacteroidota^24,39^. We also find a secondary shift associated with the evolution of mammals and birds. The dominance of Bacillota in these clades, particularly the divergence of Bacillota_A (primarily Clostridiales) in mammals and Bacillota_D (primarily Lactobacillales) in birds, likely reflects the distinct anatomical and physiological requirements of the gastrointestinal tracts between the two host clades^14^. Our study also identifies the ratio of Bacillota to Pseudomonadota as a key marker that varies with host evolution. The higher Bacillota:Pseudomonadota ratio in terrestrial vertebrates may serve as a hallmark of a gut environment that is biologically filtered and less susceptible to transient environmental influx—supported by work documenting the environmental sensitivity observed in fish and insect gut microbiomes^10,40^. These patterns can be examined and retested in the future with data types such as metagenomic, metatranscriptomic, and long-read sequencing data.

### Utility of GMToL for taxon-level inference

Future studies can leverage GMToL to determine how either their host(s) of interest or microbe(s) of interest compare to the rest of the gut microbiome tree of life. Traditionally, research focusing on the gut microbiomes of a particular animal host or group of hosts has relied heavily on manual literature reviews to draw comparisons with other taxa. This process is limited by what the authors of previous studies decide to mention and lacks the benefit of standardized processing across different studies^25^. Similarly, when investigating specific gut microbes of interest, authors frequently resort to exhaustive literature searches to determine the known host range and ecological niche of a bacterium. GMToL, in contrast, particularly with its integration into Qiita ^41^, provides authors with a rapid and reproducible way of integrating their data into the rest of the animal kingdom to draw direct evolutionary inferences on their particular host. Rather than relying on qualitative comparisons from previous publications, researchers can now quantitatively place their findings within a global context^16^. As we demonstrated with *Akkermansia muciniphila, Faecalibacterium prausnitzii*, and *Fusobacterium* spp., authors can map a microbe of interest across the host tree of life to visualize its pervasiveness and identify evolutionary conservation or specialization. By shifting from static literature-based comparisons to an active, data-driven framework, GMToL enables the identification of ancestral microbial states and the detection of convergent evolution in gut communities across vastly different host lineages^29^.

### Limitations and future directions

This work demonstrates the enduring value of 16S rRNA gene sequencing for large-scale meta-analyses. Although shotgun metagenomics offers greater functional depth, the breadth of 16S data that exists across diverse hosts allows for the investigation of deep evolutionary patterns that remain invisible in smaller, more focused datasets. However, more modern technologies, such as shotgun metagenomics using long-read sequencing^42^, should be applied at scale to produce higher-resolution data given the limitations of 16S when it comes to inferring microbial function and assigning taxonomy at the species level. Higher-resolution data, combined with improved environmental controls, would allow for the delineation between biologically-relevant bacterial strains, and likely contaminants^43^. Despite these limitations, our study provides a critical baseline for assessing gut microbiota across the animal kingdom, and serves as a vital reference for future studies on animal gut microbiome ecology and evolution.

## Methods

### KEY RESOURCES TABLE

A complete Key Resources Table summarizing software, code, deposited data, and other resources used in this study is provided as Table S6.

### RESOURCE AVAILABILITY

#### Lead contact

Further information and requests for resources should be directed to and will be fulfilled by the lead contact, Rob Knight.

#### Materials availability

This study did not generate new unique reagents or biological materials.

#### Data and code availability

- All raw 16S rRNA gene sequencing data, BIOM tables, and sample metadata aggregated as part of the Gut Microbiome Tree of Life (GMToL) are publicly available through the Qiita data portal at https://qiita.ucsd.edu/GMToL/ as of the date of publication. Per-study Qiita identifiers and original ENA/SRA accession numbers are listed in Data S1. An example meta-analysis demonstrating use of the GMToL resource through Qiita’s built-in analytical tools is publicly available at https://qiita.ucsd.edu/analysis/description/95670/.
- All original code has been deposited at GitHub and is publicly available as of the date of publication. The qiimeseq tool for fetching and processing sequence data and study metadata from ENA is available at https://github.com/luisxxu/qiimeseq, and all code for downstream analyses is available at https://github.com/samd1993/GMToL. DOIs are listed in the Key Resources Table.
- Any additional information required to reanalyze the data reported in this paper is available from the lead contact upon request.

### EXPERIMENTAL MODEL AND STUDY PARTICIPANT DETAILS

This study did not involve the generation of new biological samples, animals, or human subjects. All gut microbiome data analyzed here were aggregated from previously published 16S rRNA gene amplicon sequencing studies that were publicly available at the time of data collection. The aggregated dataset comprises 17,366 samples spanning 1,553 host species, 435 families, 26 host classes, and 9 phyla within the Bilateria clade. Per-sample metadata, including host species, host class, study identifiers, and accession numbers, are provided in Data S1. Search strategy, inclusion criteria, and sample-level filtering are described under METHOD DETAILS, “Data mining and aggregation of public gut microbiome data.”

### METHOD DETAILS

#### Datamining and aggregation of public gut microbiome data

To build a representative comparative data set of gut microbiota across the animal kingdom, we used *Web of Science* to search for 16S gut microbiome studies on animals spanning the animal kingdom, with the main objective of targeting non-mammalian and non-avian hosts. We supplemented *Web of Science* search queries, with *Google Scholar* searches for particularly rare taxa, such as annelids, echinoderms, and other invertebrates, that did not show up in the initial queries. We did not include years in our search queries since 16S high throughput sequencing studies only appear after 2006, and all studies in this dataset occur after 2012. We included studies that used either V3-V4 or V4 primers and excluded studies without metadata or accession codes to their raw data. To aggregate information from the selected studies, we developed qiimeseq, a tool to fetch and process sequence data and attach curated study metadata, such as host species taxonomy (Species, Genus, Family, Order, Class, Phylum). To ensure adequate replication per host species, we subsampled a maximum of 20 gut microbiome samples per study; in total 3,900 samples were collected for single-host species studies. We refrained from using studies of hosts in captivity unless a study on captive animals was the only one available for that host. We did not include studies of juveniles, ill animals, or animals given medical treatments, particularly antibiotics. Data S1 provides metadata for each study, with host accession numbers. For single-host species studies, we aggregated information from 185 studies spanning 15 host classes, of which seven were invertebrates. Three studies included two different host species, leading to a total of 198 hosts from 185 different studies, and 172 unique host species.

We also merged all comparative gut microbiome studies on animals and combined these with the single host species studies, totaling 17,366 samples across 284 studies. We considered comparative studies as any study with 3 or more animal hosts. Similar to the single host species studies, we only included comparative studies with available metadata and accession codes to raw sequence files that pertained to 16S V4 or V3-V4 targeted microbiome samples. We excluded studies that lacked metadata to differentiate which samples belonged to which host. In total, our dataset spanned 1,553 host species, 435 families, 26 classes, 9 phyla with an average of 9 replicates per host species across the entire dataset.

#### Data processing

To process the aggregated data, we ran Deblur (v1.1.1)^44^ on a per-study basis on the forward reads with default settings, trimming at 150bp, and then merged all reads into a single count table. We built a multi-kingdom phylogenetic tree using SEPP (implemented as the q2-fragment-insertion plugin in QIIME 2 v2024.10), which allows for the insertion of multiple primer regions into a single backbone phylogeny^45^, to insert our fragments into the Greengenes 2 (v2024.9) phylogeny^27^. We only used the forward reads because a majority of studies shared the same starting forward position, either at 515F or 341F of the 16S gene in contrast to the reverse reads which varied greatly in starting position. To assign taxonomy we used a Greengenes 2 classifier trained on the entire 16S region to ensure both V3 and V4 regions were classified properly.

#### Data analysis

For analyses that did not require the complete raw data, we filtered singletons and reads that did not show up in 2 or more samples to avoid spurious reads while still trying to retain the captured bacterial diversity in this study. For analyses that required merged samples by host species, we pooled samples and used the median relative abundance in lieu of the sum relative abundance to avoid a single sample having a disproportionate effect on the pooled relative abundance. We used these relative abundances to depict Figure 2 which highlighted prominent gut microbial phyla of each host species.

To generate a phylogenetic tree of the hosts we used TimeTree 5 (timetree.org)^30^, which calculates time-calibrated phylogenies of user inputted taxa. We utilized NCBI taxonomy codes for host names that did not match the NCBI database. The resulting host tree was used for all subsequent analyses that involved host phylogeny annotations. To generate the annotated host trees, we mapped either presence/absence (>0.01%) or the relative abundance of the microbe of interest as bars at the tips of the host tree. We then used the Interactive Tree of Life web tool (iTOL v7; https://itol.embl.de/) to visualize the tip annotations across our host tree.

To predict ancestral gut microbiome compositions, we utilized ABDOMEN (v1.0), which calculates Brownian-motion derived ancestral states with microbiome compositional data as an input for each host as a “trait”^29^. For ABDOMEN parameters, we set our detection threshold to a relative abundance of 0.00001 and a prior to ‘empirical’ at 4 chains and 2,000 iterations. The resulting output provides ancestral state gut microbiome compositions at whatever taxonomic rank was inputted. We ran ABDOMEN at the phyla and order level. To focus on prominent taxa, we selected the most relatively abundant taxa of each node from the resulting ancestral states to serve as the “trait” for that node for Figure 3.

For alpha and beta diversity analyses, we rarefied all samples to 1,000 reads in order to normalize sequencing depth while retaining 16,396 of the 17,366 samples. For alpha diversity metrics we used Faith’s phylogenetic diversity^46^ and Shannon’s diversity^47^. For beta-diversity analyses, we utilized the unweighted UniFrac metric^48^. While other metrics are available, we specifically relied on UniFrac as it takes into account all microbial taxonomic ranks, and we have previously shown that focusing on a single rank for comparative animal gut microbiome studies can be misleading^16^. Statistical significance was determined by pairwise PERMANOVA test on host class and included environmental samples as outgroups. Each environmental category– algae, sediment, saline water, non-saline water–was merged together to represent the environment as its own class in all analyses given their tight clustering and resemblance to one another relative to our gut samples. To contextualize the primary axes of the Principal Coordinates Analysis (PCoA), per-sample relative abundances of Bacillota_A, Pseudomonadota, and Bacteroidota were computed by summing taxonomically assigned feature counts within each phylum and dividing by sample read depth. These abundances were used as sample weights in Gaussian kernel density estimates^49^ projected onto the PC1 and PC2 axes, enabling direct visual comparison of phylum enrichment against the ordination structure.

To generate heatmaps of microbial taxa associations across host groups we utilized the q2- feature-table heatmap plugin in QIIME 2 v2024.10^50^. To focus on prominent taxa, we filtered the ASV table to consist of only features with more than 100 reads and showing up in at least 5 samples. We then generated dendrogram trees to connect the samples and features using Euclidean distances to represent the connections between groups. We also opted for log-transformed color gradients of the relative abundances since taxa varied widely in very low abundances to very high abundances. Host-class-resolved hexbin plots were generated in matplotlib using a two-layer approach^51^: a grey background hexbin spanning all samples provided spatial context, and a log-normalized focal-class overlay emphasized the ordination footprint of each taxonomic group, with bin transparency scaled logarithmically to avoid saturation at high-density regions.

We identified differential taxa across host classes using Bayesian Inferential Regression for Differential Microbiome Analysis (BIRDMAn v0.0.4)^38^. We chose Class because we were interested in broad comparisons across the animal kingdom as opposed to differences between closely related host species. Microbial taxa were considered credibly associated with a Class if their Bayesian credible interval contained only negative values or only positive values. The former was assigned as “negatively associated” with the phenotype and the latter was assigned as “positively associated.” A log-ratio was then constructed for the phenotype as the natural logarithm of the sum of all “positively associated” taxa divided by the sum of all “negatively associated” taxa. If the sum of either positively or negatively associated taxa was zero, the sample was not assigned a ratio for that phenotype. To visualize differentially abundant taxa across vertebrates and invertebrates as a whole, we employed BIRDMAn to ascertain the top 100 genera ranked by absolute posterior mean of the vertebrate coefficient. Inference was performed using Hamiltonian Monte Carlo chains (4 chains, 500 iterations each) yielding 2,000 posterior draws per genus. For each of the top 25 vertebrate-enriched and top 25 invertebrate-enriched genera, the full posterior distribution of the Vertebrate coefficient was summarized as a kernel density estimate (bandwidth = 0.3) and displayed as a ridgeline plot, with genera ordered by posterior mean. Taxa whose 94% highest density interval excluded zero on both sides were considered credibly associated with host vertebrate status.

### QUANTIFICATION AND STATISTICAL ANALYSIS

Statistical methods are described in detail under METHOD DETAILS alongside the analytical workflows in which they were applied. Briefly, comparisons of phylum-level relative abundances across host classes and across vertebrate versus invertebrate hosts were performed with the Kruskal-Wallis H test followed by Mann-Whitney U pairwise tests; correlations between phylum relative abundance and host divergence time from Homo sapiens were assessed with Spearman’s rank correlation. Differences in beta-diversity (unweighted UniFrac) across host classes were tested by pairwise PERMANOVA (999 permutations) using QIIME 2. Differences in alpha-diversity (Faith’s phylogenetic diversity, Shannon’s index) across host classes were tested by Kruskal-Wallis H followed by pairwise Mann-Whitney U with Benjamini-Hochberg multiple-testing correction. Differential abundance was inferred with BIRDMAn (Bayesian Inferential Regression for Differential Microbiome Analysis) using 4 Hamiltonian Monte Carlo chains of 500 iterations (2,000 posterior draws). Microbial taxa were considered credibly associated with a host group when their 94% highest density posterior interval excluded zero. Ancestral state reconstructions of microbiome composition were performed with ABDOMEN v1.0 (multivariate Brownian Motion model) using 4 chains, 2,000 iterations, an empirical prior, and a relative-abundance detection threshold of 1e-5. Sample sizes (n), test statistics (H, F, r, μ), p-values, degrees of freedom, and credible intervals (CI) for each analysis are reported in the corresponding sections of the Results and figure legends, and full per-test outputs are provided in Tables S2-S5.

## Data Availability

The Gut Microbiome Tree of Life has been made into a user-friendly data portal available at https://qiita.ucsd.edu/GMToL/. Users can download raw FASTQs, BIOM tables, and sample metadata and conduct meta-analyses using Qiita’s built-in analytical tools or with their own custom tools. An example meta-analysis demonstrating use of the GMToL resource through Qiita is publicly available at https://qiita.ucsd.edu/analysis/description/95670/. All code for fetching and sequencing data and study metadata from ENA can be found at https://github.com/luisxxu/qiimeseq and all code for downstream analyses can be found at https://github.com/samd1993/GMToL.

## Acknowledgements

We would like to acknowledge the Gut Microbiome Tree of Life Project team which involved dozens of undergraduate and graduate researchers who helped collect and annotate the large amount of study metadata that needed to be standardized.

