## Supplementary Information for "An expansive animal gut microbiome dataset elucidates major compositional shifts across bilaterian evolution"

**Table S1**. Overview of large-scale comparative gut microbiome studies (>30 host species).

| **Study** | **Host** | **Sample size** | **no. of species** | **# of samples per species** | **# of host classes** | **year** |
|---|---|---|---|---|---|---|
| Muegge *et al*. (2011) | Mammals | 38 | 33 | ~1 | 1 | 2011 |
| Jonge *et al*. (2022) | Mammals | 66 | 52 | ~1 | 1 | 2022 |
| Youngblut *et al*. (2019) | Vertebrates | 213 | 128 | ~2 | 4 | 2019 |
| Milani *et al*. (2020) | Mammals | 250 | 77 | ~3 | 1 | 2020 |
| Kim *et al*. (2021) | Fish | 227 | 85 | ~3 | 1 | 2021 |
| Hird *et al*. (2015) | Birds | 129 | 46 | ~3 | 1 | 2015 |
| Minich *et al*. (2022) | Fish | 416 | 101 | ~4 | 1 | 2022 |
| Yang et al. (2022) | All animals | 2530 | 467 | ~5 | 9 | 2022 |
| Song *et al*. (2020) | Vertebrates | 6141 | 987 | ~5 | 4 | 2020 |
| Hoffbeck *et al*. (2023) | Reptiles | 745 | 91 | ~7 | 1 | 2023 |
| Ma et al. (2021) | All animals | 4903 | 318 | ~15 | 10 | 2024 |
| **Degregori *et al.* (GMToL)** | **All animals** | **17,366** | **1,553** | **~8.5** | **26** | **2026** |

**Table S2**. Kruskal-Wallis results for CLR-transformed relative abundance of 4 microbial phyla between vertebrate and invertebrate hosts

| **Phyla** | **H-stat** | **P-value** |
|---|---|---|
| Bacillota_A | 886.517 | 8.369e-195 |
| Pseudomonadota | 120.298 | 5.44e-28 |
| Bacteroidota | 58.537 | 1.994e-14 |
| Bacillota_D | 60.997 | 5.714e-15 |

**Table S3.** Kruskal-Wallis results comparing alpha diversity (Faith's PD) across host classes.

| Group 1 | Group 2 | H | p-value | q-value |
|---|---|---|---|---|
| Annelida (n=46) | Arthropoda (n=989) | 68.487 | 1.28E-16 | 9.20E-16 |
| | Chordata (n=3251) | 51.4101 | 7.50E-13 | 4.50E-12 |
| | Cnidaria (n=25) | 40.4035 | 2.07E-10 | 8.26E-10 |
| | Echinodermata (n=27) | 23.3626 | 1.34E-06 | 4.39E-06 |
| | Free-living (n=151) | 4.77798 | 2.88E-02 | 3.84E-02 |
| | Mollusca (n=215) | 49.9166 | 1.60E-12 | 8.25E-12 |
| | water (n=26) | 11.6389 | 6.46E-04 | 1.37E-03 |
| Arthropoda (n=989) | Chordata (n=3251) | 99.8468 | 1.65E-23 | 1.98E-22 |
| | Cnidaria (n=25) | 7.53823 | 6.04E-03 | 9.88E-03 |
| | Echinodermata (n=27) | 0.01848 | 8.92E-01 | 8.92E-01 |
| | Free-living (n=151) | 149.8 | 1.92E-34 | 6.90E-33 |
| | Mollusca (n=215) | 5.44249 | 1.97E-02 | 2.83E-02 |
| | water (n=26) | 18.3232 | 1.86E-05 | 4.47E-05 |
| Chordata (n=3251) | Cnidaria (n=25) | 21.5647 | 3.42E-06 | 9.47E-06 |
| | Echinodermata (n=27) | 3.3925 | 6.55E-02 | 7.86E-02 |
| | Free-living (n=151) | 104.729 | 1.40E-24 | 2.52E-23 |
| | Mollusca (n=215) | 5.26591 | 2.17E-02 | 3.01E-02 |
| | water (n=26) | 6.65174 | 9.91E-03 | 1.55E-02 |
| Cnidaria (n=25) | Echinodermata (n=27) | 3.59351 | 5.80E-02 | 7.20E-02 |
| | Free-living (n=151) | 46.5236 | 9.05E-12 | 4.07E-11 |
| | Mollusca (n=215) | 10.876 | 9.74E-04 | 1.95E-03 |
| | water (n=26) | 26.46 | 2.69E-07 | 9.69E-07 |
| Echinodermata (n=27) | Free-living (n=151) | 22.7213 | 1.87E-06 | 5.62E-06 |
| | Mollusca (n=215) | 0.55972 | 4.54E-01 | 4.96E-01 |
| | water (n=26) | 6.02898 | 1.41E-02 | 2.11E-02 |
| Free-living (n=151) | Mollusca (n=215) | 83.0658 | 7.94E-20 | 7.14E-19 |
| | water (n=26) | 8.92651 | 2.81E-03 | 5.06E-03 |
| Mollusca (n=215) | water (n=26) | 9.26576 | 2.33E-03 | 4.42E-03 |

**Table S4**. Most prevalent gut microbes across the tree of life.

| No. of host Classes observed in | Taxon |
|---|---|
| 20 | Pseudomonadota; g__Escherichia; s__ |
| 20 | Actinomycetota; g__Corynebacterium; s__Corynebacterium kefirresidentii |
| 18 | Pseudomonadota; g__Acinetobacter; s__ |
| 18 | Cyanobacteriota; g__Caldora; s__Caldora sp010672925 |
| 18 | Actinomycetota; g__Micrococcus; s__Micrococcus luteus |
| 17 | Bacillota_I; g__Lactococcus_A_346120; s__ |
| 17 | Actinomycetota; g__Lawsonella; s__ |
| 16 | Pseudomonadota; g__Enterobacter_B_713587; s__Enterobacter_B_713587 kobei |
| 16 | Bacteroidota; g__Phocaeicola_A; s__Phocaeicola_A vulgatus |
| 16 | Bacillota_I; g__Streptococcus; s__Streptococcus thermophilus |
| 16 | Pseudomonadota; g__Methylobacterium; s__Methylobacterium thiocyanatum |
| 15 | Pseudomonadota; g__Aeromonas; s__ |
| 15 | Pseudomonadota; g__Ralstonia; s__Ralstonia mannitolilytica |
| 15 | Bacillota_A_368345; g__Faecalimicrobium; s__Faecalimicrobium dakarense |
| 15 | Bacillota_A_368345; g__Paraclostridium; s__Paraclostridium sordellii |
| 15 | Bacillota_I; g__Bacillus_A; s__ |
| 15 | Pseudomonadota; g__Acinetobacter; s__Acinetobacter schindleri |
| 15 | Pseudomonadota; g__Moraxella_A; s__ |
| 15 | Pseudomonadota; g__Sphingomonas_L_486704; s__Sphingomonas_L_486704 oligophenolica |
| 15 | Pseudomonadota; g__Bradyrhizobium_503372; s__Bradyrhizobium sp000938255 |
| 15 | Pseudomonadota; g__Methylobacterium; s__Methylobacterium radiotolerans |
| 15 | Actinomycetota; g__Janibacter_A_390549; s__Janibacter_A_390549 hoylei |
| 15 | Pseudomonadota; g__Haemophilus_D_735815; s__ |
| 14 | Bacillota_A_368345; g__Sarcina_200052; s__ |
| 14 | Bacillota_I; g__Enterococcus_H_360604; s__Enterococcus_H_360604 faecalis |
| 14 | Actinomycetota; g__Cutibacterium; s__Cutibacterium acnes |
| 14 | Pseudomonadota; g__TMED48; s__TMED48 sp002591625 |
| 14 | Pseudomonadota; g__Pelomonas; s__Pelomonas saccharophila |

**Table S5.** Top 50 differentially abundant bacterial genera between vertebrate and invertebrate gut microbiomes (top 25 enriched in each host group). Posterior mean log-fold change and 95% highest-density interval (HDI) from a Bayesian negative-binomial multilevel model (BIRDMAn) with Chordata (vertebrate vs. invertebrate) as the predictor of interest and study ID as a covariate. Positive values indicate enrichment in vertebrates relative to invertebrates; negative values indicate enrichment in invertebrates. Genera are sorted by the magnitude of the posterior mean within each group. Genus and family labels follow the GTDB taxonomy used in the source feature table; trailing numeric placeholders have been removed for readability.

| Phylum | Family | Genus | Mean log-fold | 95% HDI |
|---|---|---|---|---|
| Vertebrate-associated (top 25) | | | | |
| Bacillota_I | Erysipelotrichaceae | *Amedibacillus* | +12.66 | (+9.41, +15.91) |
| Bacillota_I | Coprobacillaceae | *Beduini* | +12.54 | (+9.30, +15.67) |
| Cyanobacteriota | Gastranaerophilaceae | *UBA2813* | +12.44 | (+9.17, +15.66) |
| Bacillota_I | Erysipelotrichaceae | *Faecalitalea* | +12.20 | (+8.70, +15.54) |
| Bacillota_A | Lachnospiraceae | *Hungatella_A* | +12.08 | (+9.40, +14.77) |
| Bacteroidota | UBA932 | *Coprenecus* | +11.98 | (+9.36, +14.75) |
| Bacteroidota | Paludibacteraceae | *RF16* | +11.87 | (+8.94, +14.68) |
| Bacillota_I | Coprobacillaceae | *Stoquefichus* | +11.86 | (+9.38, +14.15) |
| Bacillota_A | Clostridiaceae | *Clostridium_H* | +11.65 | (+8.07, +14.83) |
| Verrucomicrobiota | Victivallaceae | *Victivallis* | +11.65 | (+8.84, +14.40) |
| Bacteroidota | Coprobacteraceae | *Coprobacter* | +11.62 | (+8.07, +14.98) |
| Bacillota_A | Acutalibacteraceae | *Anaeromassilibacillus* | +11.60 | (+8.80, +14.53) |
| Verrucomicrobiota | UBA1067 | *Spyradenecus* | +11.59 | (+8.20, +14.97) |
| Thermoplasmatota | Methanomethylophilaceae | *UBA71* | +11.57 | (+8.22, +15.04) |
| Actinomycetota | Eggerthellaceae | *Slackia* | +11.35 | (+9.12, +14.05) |
| Bacillota_A | Lachnospiraceae | *Suipraeoptans* | +11.30 | (+7.87, +14.96) |
| Actinomycetota | Eggerthellaceae | *Ellagibacter* | +11.29 | (+7.92, +15.25) |
| Bacillota_A | Eubacteriaceae | *Eubacterium_O* | +11.29 | (+8.41, +13.73) |
| Bacillota_A | Ruminococcaceae | *Soleaferrea* | +11.27 | (+8.04, +14.54) |
| Bacillota_A | Oscillospiraceae | *Oscillibacter* | +11.25 | (+8.18, +14.69) |
| Bacillota_A | Lachnospiraceae | *Otoolea* | +11.19 | (+8.74, +14.13) |
| Bacillota_A | Lachnospiraceae | *Sellimonas* | +11.18 | (+8.23, +14.34) |
| Bacillota_A | Lachnospiraceae | *Porcincola* | +11.18 | (+8.97, +13.41) |
| Bacteroidota | Rikenellaceae | *Rikenella* | +11.17 | (+7.39, +14.79) |
| Bacillota_A | CAG-314 | *SFEB01* | +11.15 | (+7.68, +14.63) |
| Invertebrate-associated (top 25) | | | | |
| Pseudomonadota | Cellvibrionaceae | *Saccharophagus* | -11.53 | (-16.07, -7.10) |
| Pseudomonadota | Rhodobacteraceae | *Pseudothioclava* | -10.73 | (-15.63, -6.46) |
| Bacillota_I | Staphylococcaceae | *Mammaliicoccus* | -10.71 | (-15.38, -6.29) |
| Actinomycetota | Actinomycetaceae | *Winkia* | -10.42 | (-15.32, -5.40) |
| Pseudomonadota | Rhizobiaceae | *Roseitalea* | -10.26 | (-15.41, -5.77) |
| Actinomycetota | Pseudonocardiaceae | *Allosaccharopolyspora* | -9.81 | (-13.64, -6.07) |
| Pseudomonadota | Sphingomonadaceae | *Pacificimonas* | -9.78 | (-14.77, -5.06) |
| Pseudomonadota | Sphingomonadaceae | *Pontixanthobacter* | -9.62 | (-13.06, -6.81) |
| Pseudomonadota | Rhodobacteraceae | *Jhaorihella* | -9.56 | (-13.01, -5.99) |
| Actinomycetota | Micrococcaceae | *Auritidibacter* | -9.55 | (-15.38, -4.11) |
| Pseudomonadota | Micavibrionaceae | *UBA2137* | -9.49 | (-14.44, -4.70) |
| Bacillota_A | Lachnospiraceae | *UBA629* | -9.26 | (-14.81, -4.36) |
| Actinomycetota | Microbacteriaceae | *Pseudoclavibacter_B* | -9.18 | (-14.99, -4.40) |
| Bacteroidota | Rubricoccaceae | *Unclassified* | -9.17 | (-14.24, -4.17) |

| | | | | |
|---|---|---|---|---|
| Pseudomonadota | Caulobacteraceae | *CAHJWH01* | -9.06 | (-14.69, -4.26) |
| Pseudomonadota | DSM-100316 | *Unclassified* | -9.01 | (-13.57, -3.68) |
| Bacteroidota | Saprospiraceae | *Neolewinella* | -8.90 | (-11.56, -6.47) |
| Pseudomonadota | Vibrionaceae | *Vibrio_A* | -8.79 | (-10.70, -6.55) |
| Pseudomonadota | Salinisphaeraceae | *Salinisphaera* | -8.72 | (-11.63, -6.09) |
| Bacillota_A | Vallitaleaceae_A | *Vallitalea* | -8.66 | (-14.23, -3.84) |
| Bacillota_I | Anoxybacillaceae | *Anoxybacillus_A* | -8.57 | (-10.26, -6.85) |
| Bacteroidota | Tannerellaceae | *Tannerella* | -8.46 | (-11.25, -5.38) |
| Pseudomonadota | Enterobacteriaceae_A | *Mangrovibacter* | -8.40 | (-15.16, -2.14) |
| Pseudomonadota | Maricaulaceae | *Hellea* | -8.37 | (-10.61, -6.46) |
| Planctomycetota | Pirellulaceae | *Stieleria* | -8.37 | (-11.18, -5.72) |

### Supplementary figures

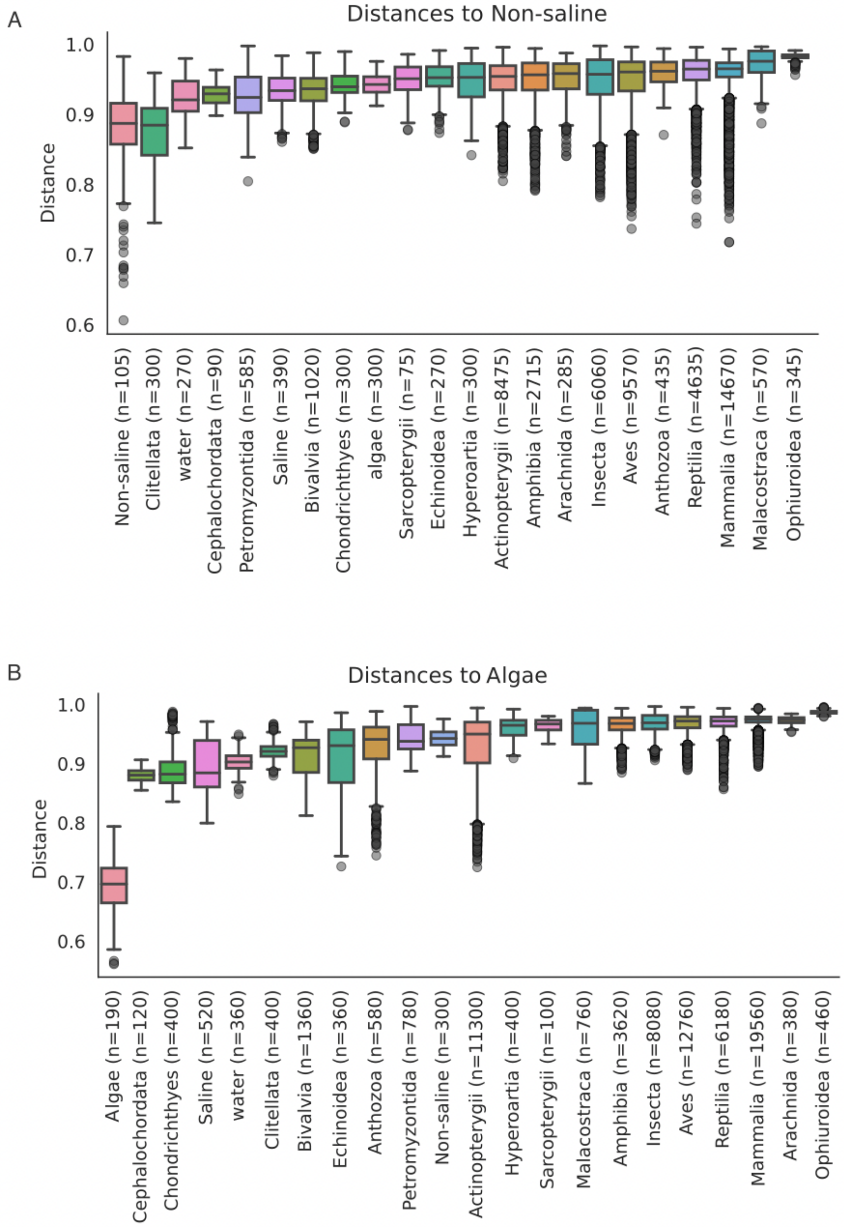


Fig S1. Unweighted UniFrac distances relative to (a) non-saline environmental samples and (b) marine algae samples. Each bar represents samples of a host Class or environmental sample type.

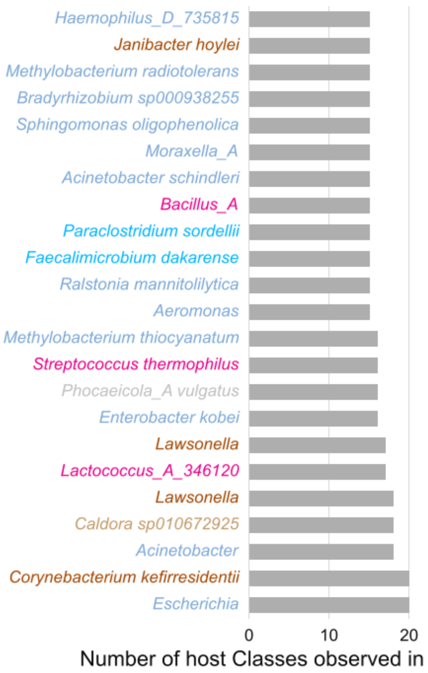


Figure S2. The most prevalent gut microbes (present in at least 9 host classes) across the entire dataset shown at the species level or next best classification rank.

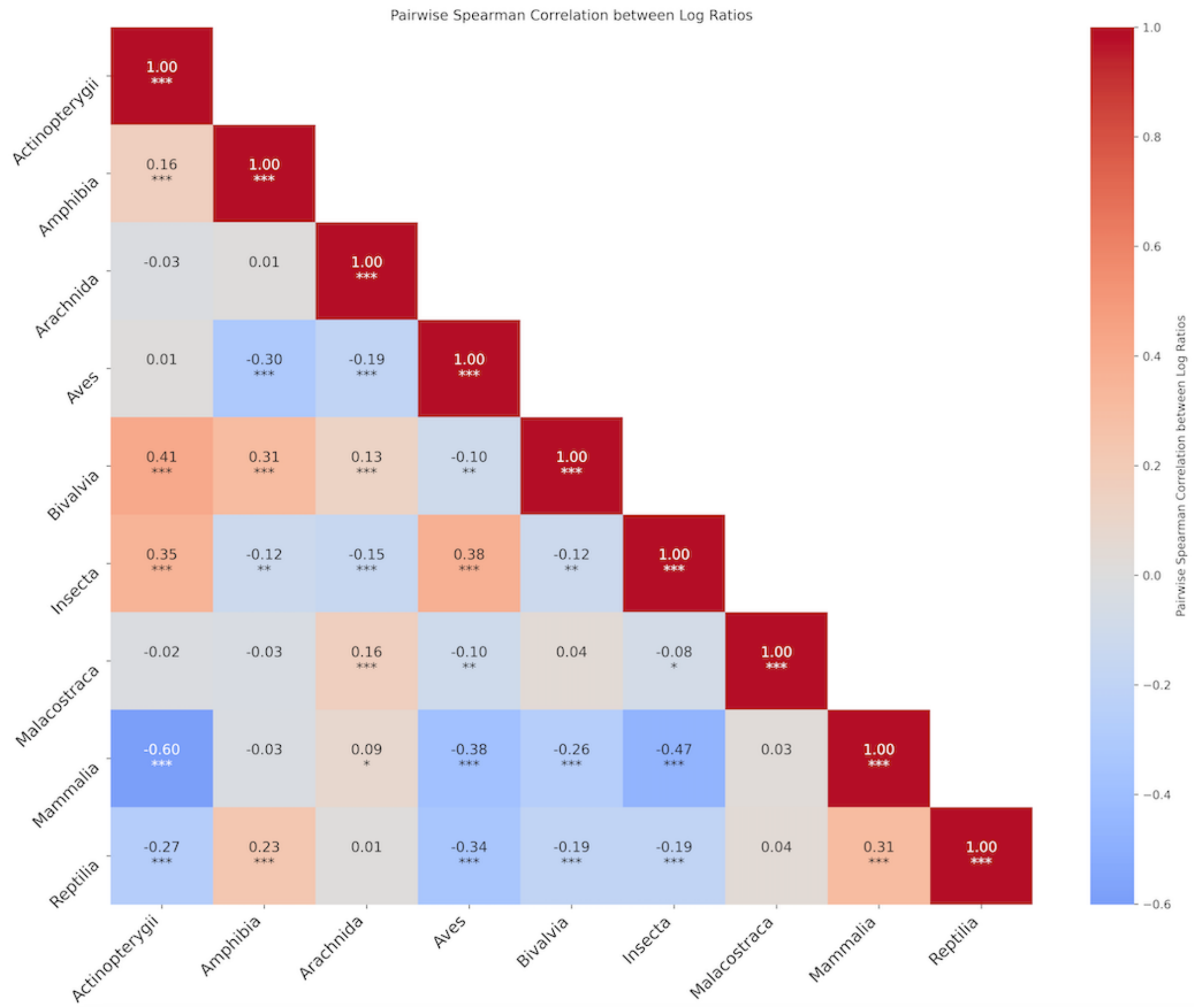


Fig S3. A Spearman correlation heatmap of host classes based off the log ratios of enriched versus depleted taxa from our BIRDMAn results. ***highly significant ** significant *notable.
